# BioEngine: scalable execution and adaptation of bioimage AI through agent-readable interfaces

**DOI:** 10.64898/2026.04.19.719496

**Authors:** Nils Mechtel, Hugo Dettner Källander, Songtao Cheng, Hanzhao Zhang, AI4Life Horizon Europe Program Consortium, Wei Ouyang

**Affiliations:** Department of Applied Physics, Science for Life Laboratory, KTH Royal Institute of Technology, Stockholm, Sweden; AI4Life Horizon Europe Program Consortium (https://ai4life.eurobioimaging.eu/)

## Abstract

Foundation models and curated repositories have transformed bioimage AI, yet most biologists cannot readily run, adapt, or extend them on available hardware. BioEngine lls this gap as the execution and adaptation layer between curated AI and scalable compute, deployable on a laptop, workstation, or cluster. Scientists then screen models, ne-tune from the browser, enable real-time smart microscopy, and deploy analysis applications, all by describing their goal to an AI agent.

## INTRODUCTION

Public repositories and foundation models have substantially expanded the menu of bioimage AI: the BioImage Model Zoo^1^ hosts hundreds of curated, FAIR-validated models, accessible training platforms^6,7^ lower the barrier to custom workflows, and general-purpose foundation models approach universal cell segmentation across modalities and organisms^4,8^. Smart microscopy systems are closing the acquisition-to-analysis loop in real time through live ML inference^17,18^. Yet most biologists still cannot readily test, adapt, or deploy these resources on their own hardware.

This disconnect has two reinforcing causes. On the interface side, AI assistants such as the BioImage.IO Chatbot^9^ and local tools such as Omega^15^ help users navigate models and interact with images, but remain advisory or tightly scoped tools that cannot orchestrate scalable GPU workflows on behalf of the user. On the infrastructure side, GPU hardware is widely available, from workstations in individual labs to clusters in core facilities, yet configuring it for AI model execution requires specialist programming that most biologists lack. Adapting foundation models such as Cellpose-SAM^4^ and μSAM^8^ to new imaging conditions requires local training expertise. Building custom analysis pipelines that compose multiple models with downstream measurements and user interfaces requires software engineering skills that lie beyond the scope of most life science training. Research prototypes now show that agents can autonomously design and train deep-learning models^13^. The missing piece is a production layer that makes those agent capabilities accessible on the hardware researchers already have.

We developed BioEngine to bridge this disconnect (Fig. 1a). BioEngine is an open-source platform that manages AI model execution, fine-tuning, and application deployment on any GPU hardware, from a single laptop or workstation to a multi-node cluster^3^, with a one-time setup. Scientists then access the full power of that hardware through an AI agent. BioEngine’s SKILL.md contracts translate natural-language requests into GPU jobs, handling model selection, fine-tuning, and pipeline deployment with no programming required. Whoever manages the hardware controls the infrastructure. The science is in the biologist’s hands.

**Fig. 1:**
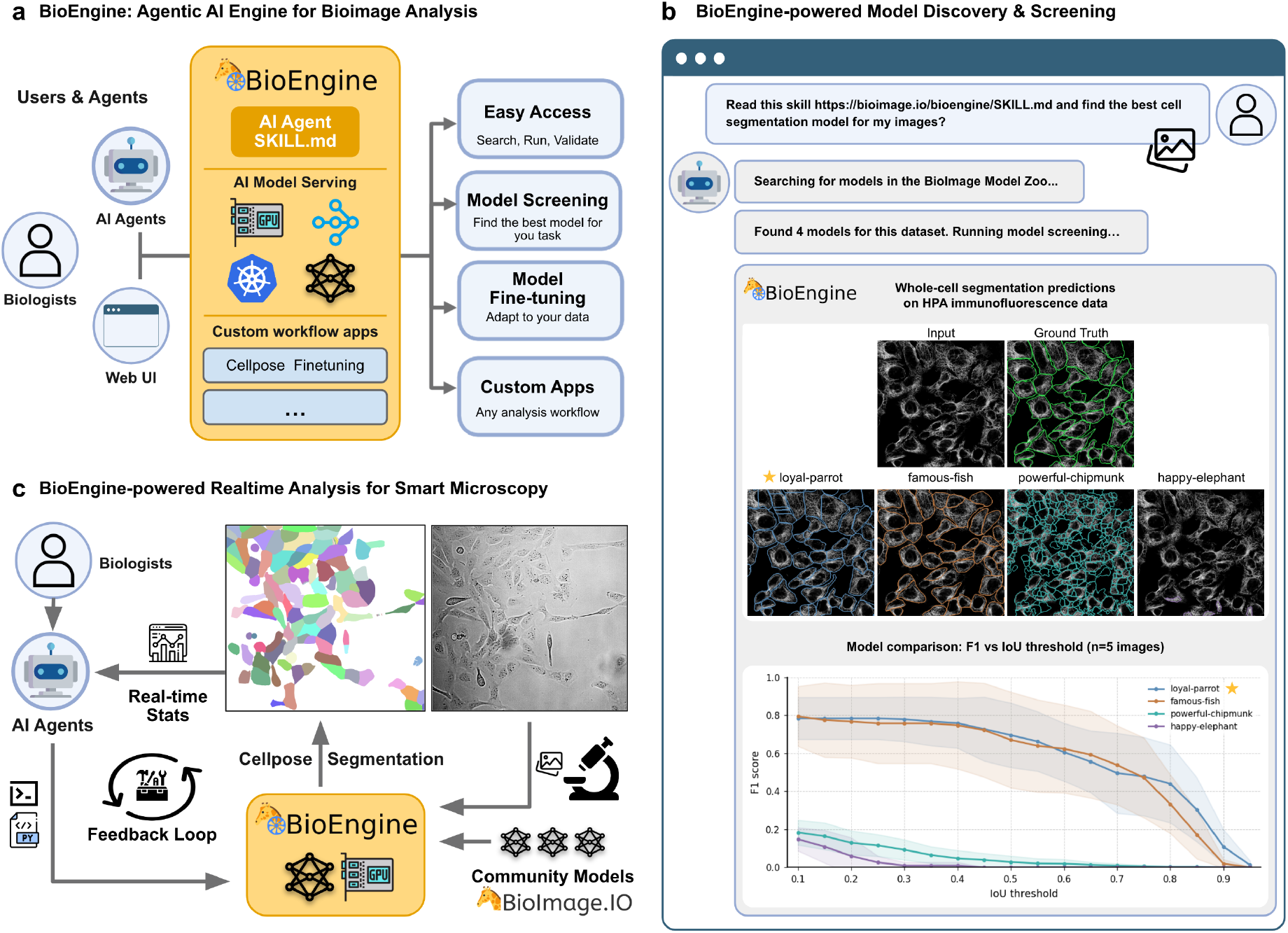
BioEngine: an agentic AI engine for bioimage analysis. a, BioEngine acts as a shared execution layer on GPU hardware, routing requests from biologists, AI agents, and web interfaces through an a services, enabling model access, screening, fine-tuning, and applicatio powered model discovery and screening. An AI agent searched 58 Bio and ranked them by mAP (mean average precision, IoU 0.1–0.95^15^, n ^14^ are shown. Loyal-parrot, trained on HPA data, ranked highest. c, Bio streamed from a microscope are segmented in real time by BioEngine feedback-guided imaging and adaptive acquisition decisions. a, BioEngine acts as a shared execution layer on GPU hardware, routing gent-readable SKILL.md contract to model inference and workflow n deployment without local software installation. b, BioEngine-Image Model Zoo candidates, filtered to 4 domain-compatible models, = 5 images). F1 vs IoU threshold curves and segmentation overlays Engine-powered real-time analysis for smart microscopy. Live images, and per-frame outputs returned to the user or AI agent enable

## RESULTS

### BioEngine: an agentic AI engine for bioimage analysis

BioEngine exposes its capabilities through a SKILL.md contract (Fig. 1a), a plain-text file format designed for general-purpose AI agents to acquire domain knowledge and invoke services directly^19^. The document lists every available analysis service with its inputs, outputs, and usage examples. A scientist describes their imaging goal in plain language to any AI agent. The agent parses the contract, selects the appropriate service, and dispatches the corresponding GPU workflow to the available hardware. Results return as segmented images, ranked comparison tables, or a live web application, with no command-line access, software installation, or IT ticket required. The following sections demonstrate four use cases served by this interface: automated model screening, real-time inference for live microscopy, browser-based collaborative fine-tuning, and deployment of agent-built analysis applications.

### Agent-driven model screening

Choosing the right AI model for a new dataset is typically a slow and manual process. A biologist must identify candidates in the literature, configure each software environment, run inference, and compare outputs by eye, a process that can take days or fail to happen at all when no single environment supports all candidates. BioEngine transforms this into an automated ranking workflow. Its model execution service runs any BioImage Model Zoo^1^ model on GPU hardware, enabling systematic comparison across many candidates within a single session. An AI agent equipped with the BioEngine SKILL.md autonomously queries the Zoo for relevant candidates, filters by documented compatibility with the input data type, runs inference on the biologist’s own images, and ranks results by mAP (mean average precision over IoU thresholds 0.1–0.95^15^). In a whole-cell segmentation task on Human Protein Atlas^10^ immunofluorescence images (microtubule channel), the agent searched 58 candidate models, excluded 54 after domain-compatibility review, ran inference for the four compatible models, and ranked them by mAP (Fig. 1b). Loyal-parrot^1^, a model specifically trained on HPA data, ranked highest, demonstrating that domain match matters and that an agent can identify this without the biologist testing each model manually.

### Real-time AI inference for agent-guided smart microscopy

The same GPU execution infrastructure used for model screening operates with sub-second latency on streaming inputs, extending its use directly to live imaging settings (Fig. 1c). When inference latency reaches this range, AI model execution becomes a real-time sensor that can close the loop between a live microscope and an AI agent. Smart microscopy platforms have demonstrated the power of this loop, from adaptive lattice light-sheet systems that detect rare cellular events and switch autonomously to high-resolution 3D acquisition^17^, to electron microscopy workflows that allocate beam time in real time based on ML predictions to achieve sevenfold acceleration^18^. What has limited broader adoption is not the microscope hardware nor the AI models themselves, but execution: running a BioImage Model Zoo model on a GPU cluster previously required specialist programming and introduced latency incompatible with live feedback.

BioEngine removes both barriers. Live images stream from the microscope to BioEngine, which schedules GPU inference and returns per-frame statistics (cell count, segmentation masks, morphological measurements, fluorescence intensities) to the controlling AI agent. The agent can then act on these outputs: adjusting focus or illumination, triggering a high-resolution volume scan on detection of a rare event, switching imaging modality, or, in systems with fluidic control, modifying the cellular environment. Because BioEngine exposes this capability through the same SKILL.md interface used for model screening and fine-tuning, any AI agent that can screen models offline can also direct real-time imaging decisions, with no additional programming. The same GPU hardware that supports batch analysis during the day is therefore capable of powering fully autonomous, hypothesis-driven imaging while samples are live on stage.

### Collaborative ne-tuning and community model contribution

Foundation models such as Cellpose-SAM^4^ and μSAM^8^ have made impressive strides towards universality, but even the most capable models fail on imaging conditions outside their training distribution. Adapting them traditionally requires a local GPU, training scripts, and days of iterative validation, steps that put fine-tuning out of reach for most biologists. BioEngine addresses this through a browser-based collaborative annotation and fine-tuning workflow that requires no local software and scales to annotators worldwide (Fig. 2a–c).

**Fig. 2:**
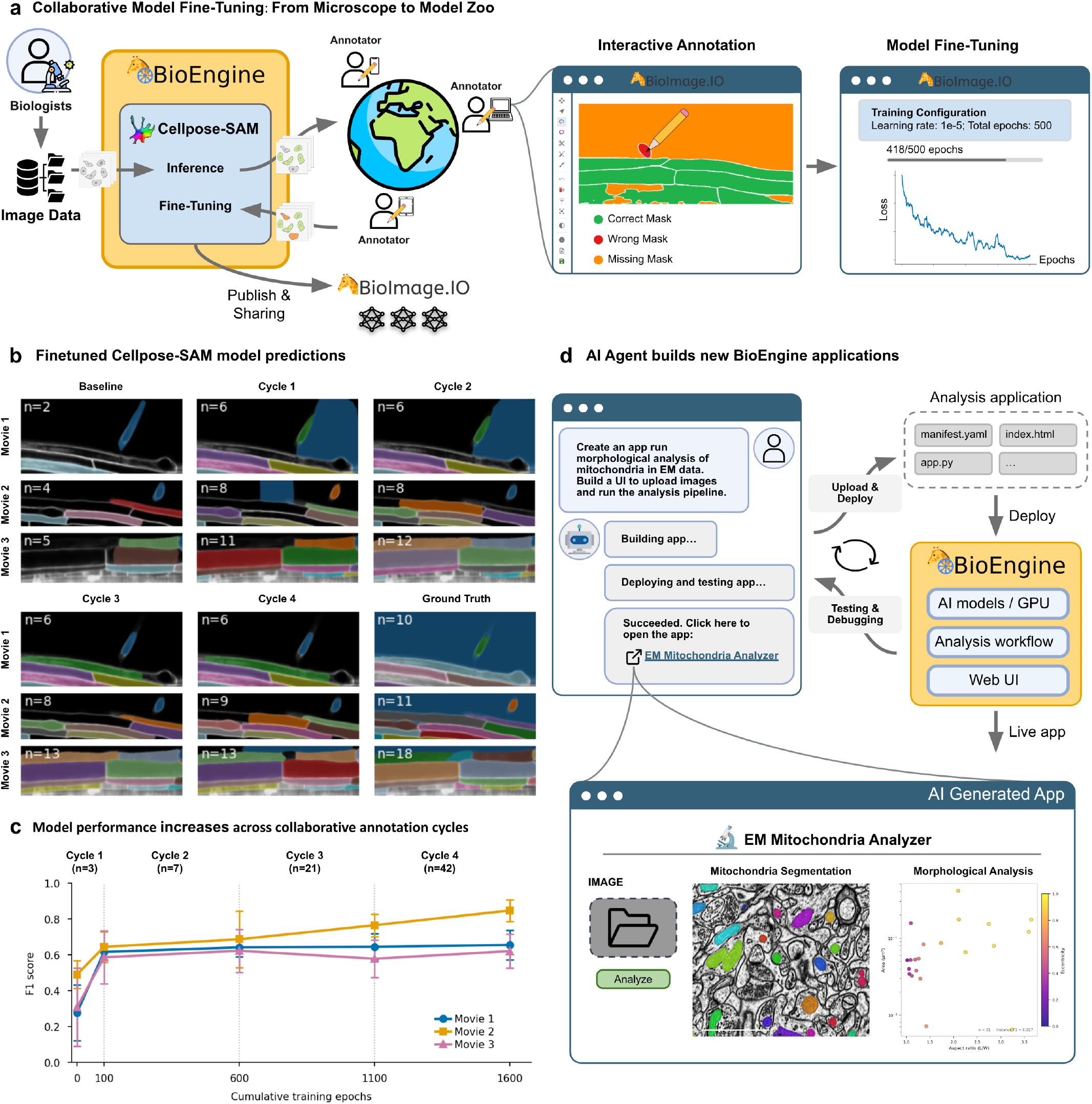
Extending BioEngine through collaborative model ne-tuning and agent-built applications. a, Collaborative fine-tuning workflow. A biologist submits microscopy images to BioEngine, which runs Cellpose-SAM^4^ to produce initial segmentations. Annotators worldwide correct those masks through a browser-based tool at BioImage.io, and each annotation cycle adds new training images and triggers GPU fine-tuning on BioEngine with one click and no local installation. The progressively improved model is published to BioImage.IO and becomes immediately available through any BioEngine deployment, extending the shared model pool for the wider community. b, Segmentation montage showing Cellpose-SAM predictions on PlantSeg^5^ *Arabidopsis* lateral root nuclei (Movies 1–3) from baseline through four collaborative fine-tuning cycles, alongside ground-truth. Baseline Cellpose-SAM fails to detect most large plant nuclei. As more annotated slices accumulate across cycles, the model progressively recovers boundaries and separates adjacent objects. c, F1 score (IoU ≥ 0.5) on held-out test slices (± s.d., n = 3 slices per movie) increases consistently across all three movies with each annotation cycle, rising from a mean of 0.36 at baseline to 0.71 after four cycles (1,600 cumulative training epochs). d, An AI agent builds new BioEngine applications by generating the analysis workflow, deployment manifest, and web interface from a single plain-language prompt. The deployed EM Mitochondria Analyzer accepts EM image uploads and returns instance segmentations with morphological profiles on the Lucchi++ FIB-SEM benchmark^11^ (neural tissue, 5 nm/px), illustrating how BioEngine can be rapidly extended with new analysis capabilities without manual programming.

The workflow centres on an interactive annotation interface hosted at BioImage.io that uses BioEngine as its GPU back end. Rather than annotating from scratch, biologists start from Cellpose-SAM pre-segmentations generated by BioEngine, correcting errors directly in the browser. Training progress is visible in real time through the same interface. Crucially, when using a shared BioEngine deployment, annotators anywhere in the world can contribute to the same session simultaneously, enabling the kind of collaborative curation that would be impossible with local software. When sufficient corrections have accumulated, fine-tuning is triggered with a single click. The improved model is automatically exported in BioImage.IO-compatible format and published to the BioImage Model Zoo, making it immediately available through any BioEngine deployment and contributing directly to the community’s shared model pool.

To demonstrate this workflow, we used the PlantSeg lateral root dataset^5^ (*Arabidopsis thaliana*, xylem pole pericycle nuclei, three confocal movies). Cellpose-SAM severely undersegments these large plant nuclei (Fig. 2b, baseline): the p90 nucleus diameter in this dataset (293– 399 px) lies well outside the model’s training range (7.5– 120 px), making it a demanding test of what collaborative fine-tuning can recover. Using existing ground-truth segmentations as proxy annotations in a simulated session, we ran four progressive annotation cycles, expanding from three slices of one movie to 42 slices across all three. With each cycle, the model received more training images and was fine-tuned for an additional 100– 500 epochs on BioEngine. Segmentation quality improved rapidly and consistently: mean F1 rose from 0.36 to 0.71 across nine held-out test slices over 1,600 cumulative training epochs (Fig. 2b–c). Gains were consistent across all three movies, demonstrating that adding data from diverse imaging conditions drives generalisation rather than overfitting. The same infrastructure, connected to a live imaging system, would allow a model to be fine-tuned and re-deployed while the sample is still on stage, eliminating the offline processing bottleneck that currently separates smart microscopy acquisition from model adaptation^12^.

### Extending BioEngine with agent-built applications

BioEngine is designed as an extensible platform. Applications are self-contained, deployable units that plug into the same GPU infrastructure and SKILL.md interface used for model screening and fine-tuning. An application can compose multiple AI models into a multi-step workflow, wrap an existing model with custom pre- and post-processing, expose an arbitrary web user interface for upload, annotation, or interactive visualisation, or combine all of the above. Because applications register themselves as services upon deployment, they become immediately discoverable and callable by AI agents through the same capability-contract mechanism. A biologist can extend BioEngine with entirely new analysis workflows tailored to their biological question, all without writing code.

To demonstrate this, an AI agent was given the BioEngine SKILL.md and a single plain-language prompt describing the desired analysis (Fig. 2d). The agent generated a deployment manifest, a GPU workflow script composing the existing model-runner service with custom analysis logic, and a web user interface, then uploaded the package to the Hypha artifact store and triggered deployment. The result, the EM Mitochondria Analyzer, is a live web application that accepts electron microscopy image uploads and returns GPU-accelerated mitochondria instance segmentation alongside a per-organelle morphological profile, all without any manual programming. The application delegates GPU inference to the model-runner service running the poisonous-spider model (a BioImage.IO Attention U-Net trained on the Lucchi++ benchmark^11^), adds watershed-based instance separation and per-instance shape analysis, and classifies mitochondria as round or elongated based on aspect ratio. Applied to the Lucchi++ FIB-SEM benchmark (neural tissue, 5 nm/px), the pipeline automatically distinguished round from elongated mitochondria and produced quantitative morphological profiles. Independent validation on 10 test slices confirmed mean instance F1 = 0.920 ± 0.037 at IoU ≥ 0.5 (Supplementary Fig. S2). The same pattern, a biologist describes a need, an agent builds and deploys the application, extends to any analytical workflow that can be expressed as a composition of GPU models and a web interface.

## DISCUSSION

BioEngine addresses a structural gap in the bioimage analysis ecosystem. Curated model repositories and foundation models have raised the ceiling of what is computationally achievable in biological imaging, but the floor, practical access to GPU execution for the models that exist, has not kept pace. BioEngine fills this gap as the execution and adaptation layer between the model and the hardware. Whether deployed on a lab workstation or an institutional cluster, it requires only a one-time setup. Scientists then access every capability through any AI agent by pasting a single SKILL.md link. The user does not need to know how BioEngine works internally. They describe what they want and receive the result. Infrastructure management stays with whoever runs the hardware. Scientific focus returns to the biologist.

The demonstrations here trace the full arc of how this layer is used. Model screening removes the days-long process of testing candidate models and replaces it with a ranked comparison generated from the biologist’s own images. Real-time inference closes the feedback loop between a live microscope and an AI agent, enabling the adaptive acquisition that smart microscopy platforms have demonstrated^17,18^ but that previously required specialist streaming infrastructure. Collaborative fine-tuning lets biologists adapt foundation models^4,8^ to new imaging conditions without a local GPU or training code, and each fine-tuned model is published back to the BioImage Model Zoo^1^ for the entire community. Agent-built applications extend BioEngine itself, allowing new composed workflows and web interfaces to be deployed by any biologist who can describe what they need. Each of these was previously impractical for most labs. Perspectives on multimodal AI in bioimaging envision exactly this kind of end-to-end, agent-driven capability as the future of the field^16^.

A broader consequence of this design is a new mode of computational biology, one in which users with no programming background interact with GPU-backed analysis systems entirely through natural language. The biologist articulates intent, the agent constructs and executes the workflow, and results arrive without the biologist ever inspecting code or configuring software. This paradigm, which we term vibe analysis by analogy with vibe coding, the emerging practice of producing software through natural language prompts without programming expertise, substantially lowers the entry barrier to bioimage AI. It also introduces a critical risk: language models can generate workflows that produce plausible-looking outputs while performing incorrect analysis, and in a setting where the biologist did not write the code and cannot readily inspect it, such errors can propagate undetected through an entire study.

Several strategies reduce this risk without requiring biologists to become programmers. Requesting intermediate outputs gives the biologist a visual checkpoint at each analysis stage without requiring code inspection. Reviewing segmentation masks before accepting cell count summaries, or per-instance measurements before aggregate statistics, lets the biologist verify that the underlying logic is correct at the pixel level. Standardised validation steps built into SKILL.md contracts, such as running a model on a known reference sample before applying it to experimental data, provide factual anchors that partially counteract hallu inated outputs. For applications intended for routine facility use, expert code review of the agent-generated application remains the most reliable safeguard. Training biologists in this new paradigm, focused not on writing code but on workflow verification, intermediate output inspection, and risk-aware prompting, is likely to become as important as traditional scripting skills in bioinformatics education. As agents become more capable and community-contributed skills accumulate, the range of well-tested, grounded capabilities available through BioEngine will grow, progressively narrowing the space in which hallucination can cause harm.

## SUPPLEMENTARY FIGURES

**Supplementary Fig. S1** | **Nucleus size distributions in PlantSeg lateral root movies motivate per-movie diameter normalisation**. Histograms of equivalent nucleus diameter (log scale) for the three PlantSeg movies (Movie 1–3) used in the collaborative fine-tuning experiment. The grey region indicates the diameter range (7.5–120 px) on which Cellpose-SAM was trained. The dashed line marks the 90th-percentile (p90) diameter per movie (M1: 399 px, M2: 355 px, M3: 293 px). All three movies lie substantially outside the Cellpose-SAM training range, requiring per-movie rescaling to bring nuclei within the model’s operating diameter window. Per-movie rescaling factors (M1 × 0.30, M2 × 0.34, M3 × 0.41) map p90 diameters to 120 px and were applied consistently to both training and test images.

**Supplementary Fig. S2** | **Quantitative validation of the EM Mitochondria Analyzer on the Lucchi++ benchmark**. Instance F1 at IoU ≥ 0.5 for the BioEngine-deployed poisonous-spider pipeline across 10 evenly spaced test slices of the Lucchi++ FIB-SEM dataset (neural tissue, 5 nm/px). The dashed line indicates the mean and the shaded band shows ± s.d. Mean F1 = 0.920 ± 0.037 (n = 10 slices).

## DATA AVAILABILITY

PlantSeg lateral root data are publicly available via the PlantSeg repository. Human Protein Atlas images and ground-truth annotations for model screening were obtained from the HPA Cell Image Segmentation Dataset^14^ (https://www.proteinatlas.org). FIB-SEM data are from the Lucchi++ benchmark^11^.

### CODE AVAILABILITY

BioEngine source code: https://github.com/aicell-lab/bioengine-worker. BioEngine SKILL.md: https://bioimage.io/skills/bioengine/SKILL.md. BioImage Colab collaborative annotation tool and BioEngine web interface: https://github.com/bioimage-io/bioimage.io. Production service: https://hypha.aicell.io (workspace: bioimage-io).

## ACKNOWLEDGEMENTS

We thank the AI4Life Horizon Europe Program Consortium members for their efforts in creating and improving the BioImage Model Zoo. We are grateful to F. Beuttenmueller for developing, improving, and maintaining the BioImage Model Zoo core specification, which is central to BioEngine’s model execution layer. We thank the participants of the BioEngine hackathon “Web and Cloud Infrastructure for AI-Powered BioImage Analysis” (Stockholm, 2023; Craig Russell, Carlos Garcia, Caterina Fuster-Barceló, Curtis Rueden, Jonas Klotz, Matthew McCormick, Nodar Gogoberidze, Benjamin Wilhelm, Andréa Papaleo, Simon Franchini, Tomaz Fogaça Vieira, Sebastian Soyer, Weize Xu, Iván Hidalgo, and Vera Galinova) for discussion, testing, and feedback on the infrastructure. We thank Curtis Rueden and Kevin W. Eliceiri for discussion and suggestions for improving BioEngine, Gisele Miranda for testing and feedback, and Emma Lundberg for discussion and support during development. We thank Jean-Yves Tinevez, Jean-Christophe Olivo-Marin, Maximilian Schulz, and John Eriksson for discussion and feedback. We thank Dorothea Dörr for administrative support in AI4Life. The authors used Claude (Anthropic) and ChatGPT (OpenAI) for assistance with coding, experiment design, and drafting of this manuscript.

This work was supported by the SciLifeLab & Wallenberg Data Driven Life Science Program (grant: KAW 2020.0239), the Göran Gustafsson Prize (grant: 2317) awarded to W.O., and the European Union’s Horizon Europe research and innovation programme under grant agreement No. 101057970 (AI4Life), awarded to W.O., and the European Union grant agreement No. 101188168 (RI-SCALE), awarded to W.O. GPU computation was enabled by the Berzelius resource provided by the Knut and Alice Wallenberg Foundation at the National Supercomputer Centre. This work is also supported by the de.NBI Cloud within the German Network for Bioinformatics Infrastructure (de.NBI) and ELIXIR-DE (Forschungszentrum Jülich and W-de.NBI-001, W-de.NBI-004, W-de.NBI-008, W-de.NBI-010, W-de.NBI-013, W-de.NBI-014, W-de.NBI-016, W-de.NBI-022).

## CONSORTIA

### AI4Life Horizon Europe Program Consortium

Anna Kreshuk, Arrate Muñoz-Barrutia, Beatriz Serrano-Solano, Caterina Fuster Barceló, Constantin Pape, Craig T. Russell, Dominik Kutra, Emma Lundberg, Estibaliz Gómez-de-Mariscal, Florian Jug, Fynn Beuttenmueller, Iván Hidalgo-Cenalmor, Jeremy Mertz, Joanna Hård, Joran Deschamps, Joshua Talks, Mariana G. Ferreira, Marcus Andersson, Matthew Hartley, Mehdi Seifi, Ricardo Henriques, Teresa Zulueta-Coarasa, Vera Galinova, Wei Ouyang, and others (full membership at https://ai4life.eurobioimaging.eu/).

## AUTHOR CONTRIBUTIONS

W.O. conceptualisation, conceived and supervised the project. N.M. and H.D.K. developed the BioEngine infrastructure and SKILL.md interfaces. S.C. and H.Z. performed experiments and evaluations. All authors contributed to writing.

## Methods

### BIOENGINE DISTRIBUTED ARCHITECTURE

BioEngine is a distributed computing platform that makes bioimage AI models and analysis workflows accessible as callable services from any network location. It is organised across three main components: a central worker process that coordinates local computation, a connectivity layer that manages connections from remote users and agents, and a Ray parallel computing cluster (v2.33.0) that executes AI models and custom applications. The worker connects to a Hypha management server on start-up to register its capabilities as accessible services, and supports three deployment modes: `slurm` mode for high-performance computing (HPC) environments using Apptainer containerised execution, `single-machine` mode for local deployment and testing, and `external-cluster` mode for attachment to an existing Ray cluster.

The worker coordinates two internal subsystems. The first monitors and manages the Ray computing cluster, collecting real-time CPU, GPU, and memory utilisation per machine alongside pending task counts with minimal overhead. The second handles deployment of AI analysis applications using Ray Serve: each application is stored as a versioned package in Hypha, with a configuration file (manifest) specifying its computational components, required software dependencies, and the list of authorised users. BioEngine retrieves each package, installs its dependencies into an isolated software environment, and launches it as a running service, allowing applications to be versioned, shared, and updated without modifying the core infrastructure.

The worker exposes management services for remote administration: real-time health checks, resource usage monitoring, controlled shutdown, and remote log access. Access control uses token-based authentication (JSON Web Tokens) with two permission levels: administrators can stop the worker, upload new applications, and run custom Python scripts on the cluster; any authenticated user can query cluster status and application availability. Because the scripting interface runs without filesystem-level isolation, it is intentionally restricted to administrators.

The reference deployment used in this study ran on a Kubernetes cluster (a widely used cloud computing platform) managed by the KubeRay operator (v1.1.1). The Ray cluster consisted of a CPU-only coordinator node (4 cores, 10 GB RAM) and an auto-scaling pool of 1–8 GPU worker nodes (8 cores, 32 GB RAM each) equipped with NVIDIA A40 GPUs (16 GB vRAM per GPU, CUDA 11.8). Node count scaled automatically with demand. A 100 GiB shared storage volume was accessible from all nodes. All communication between components used TLS-encrypted connections.

### REMOTE ACCESS AND SERVICE CONNECTIVITY

Hypha is the connectivity layer that links BioEngine workers, data storage, and client applications. It establishes persistent encrypted connections (using the WebSocket protocol) that allow AI agents, web browsers, and analysis scripts to call BioEngine services with low latency from any network, without requiring open network ports or VPN access. Hypha maintains a directory of all registered services, enabling clients to discover and route requests to the correct service automatically. Authentication uses JSON Web Tokens with role-based access control.

Hypha organises resources into two categories. Artifacts are versioned, stored packages (application code, model weights, or datasets) held in S3-compatible cloud storage and identified by a workspace-scoped name (browsable at https://hypha.aicell.io/<workspace>/artifacts/<name>).Services are live, callable analysis endpoints registered by running applications (accessible at https://hypha.aicell.io/<workspace>/services/<name>).

BioEngine applications follow a two-stage lifecycle: the application code is first stored as a versioned artifact, then activated on the Ray cluster, at which point it registers itself as a live service and becomes available to agents and users. In this study, Hypha coordinated all interactions between BioEngine workers, storage, and client applications including AI coding agents and web browser interfaces. Human-facing web applications use the same JavaScript client library as AI agents, ensuring biologists and agents access identical capabilities through their respective interfaces.

### AGENT-READABLE SKILL CONTRACTS

BioEngine exposes a SKILL.md document that describes all capabilities available on a deployed BioEngine instance. The document defines every callable operation alongside its required inputs, expected outputs, and usage examples. In practice, an agent receives the SKILL.md link, reads the contract, and plans calls to the corresponding BioEngine services. This pattern provides a stable, human-readable interface for AI agents while ensuring reliable, reproducible execution on BioEngine infrastructure. A BioEngine command-line tool, bundled with the SKILL.md, wraps all underlying service calls and provides a single entry point for both agent-driven and programmatic use, so that agents never need to construct low-level API calls directly.

### BIOIMAGE.IO MODEL EXECUTION SERVICE

The Model Runner application (v1.0.0) standardises execution of BioImage.IO models using a two-tier design that separates request handling from GPU inference. The first tier runs on CPU only and handles request routing, input validation, and model weight caching. It maintains a local cache of up to ten model packages using atomic file operations to coordinate concurrent downloads safely, is provisioned with 1 CPU, 0 GPUs, and 4 GB RAM, and limits the queue to 30 concurrent requests to prevent overload. The second tier is a GPU execution environment pre-configured with the bioimageio.core inference library (v0.10.0) and the major deep-learning frameworks (PyTorch 2.5.1 and torchvision 0.20.1, TensorFlow 2.16.1, and ONNX Runtime 1.20.1). Each GPU replica is provisioned with 1 CPU, 1 GPU, and 12 GB RAM, and the deployment scales automatically between one and two replicas based on request load. The service is accessible at https://hypha.aicell.io/bioimage-io/services/model-runner and via a Model Context Protocol (MCP) endpoint.at https://hypha.aicell.io/bioimage-io/mcp/model-runner

### CELLPOSE-SAM FINE-TUNING SERVICE

The Cellpose Fine-Tuning application (v0.0.26) provides a dedicated service for iterative, browser-initiated fine-tuning of the Cellpose-SAM model (Cellpose v4.0.7, cpsam checkpoint). It is implemented as a single Ray Serve deployment supporting iterative training cycles. Internally, a PyTorch module wraps the Transformer-based Cellpose-SAM architecture to make it compatible with the BioImage.IO model standard. The deployment is provisioned with 1 GPU, 4 CPUs, and 12 GB RAM per replica, and is limited to a single replica to prevent GPU memory conflicts across concurrent training sessions. Fine-tuning begins from the pre-trained `cpsam` checkpoint; training progress is streamed in real time to the annotation interface via callback functions, and completed models are automatically exported in BioImage.IO format with metadata and compressed weights. The service is accessible at https://hypha.aicell.io/bioimage-io/services/cellpose-finetuning.

### MODEL SCREENING EVALUATION

#### Dataset

The HPA Cell Image Segmentation Dataset^14^ (Kaimal & Thul, Zenodo; 159 annotated test images, 512 × 512 px, four-channel HPA immunofluorescence (microtubules, ER, nuclei, protein of interest)) was used to evaluate whole-cell segmentation model performance. Ground-truth cell boundaries were provided as polygon annotations generated by expert annotators. Five test images from this dataset were used for evaluation.

#### Agent screening workflow

The agent received the BioEngine SKILL.md and the plain-language task: *“Find the best whole-cell segmentation model for my HPA immunofluorescence images*.*”* It searched the BioImage Model Zoo via the BioImage.IO model execution service using nine task-relevant keyword queries (`cell segmentation`, `cellpose`,`fluorescence segmentation`,`stardist`, `2D instance segmentation`, `whole-cell`, `nucleus segmentation`, `HPA`, and `cyto`), returning 58 candidate models in total. For each candidate, the agent retrieved the model documentation to inspect training domain, required input channels, and known limitations. Models with incompatible domains were excluded without running inference: 54 models were discarded for reasons including nucleus-only segmentation targets, phase-contrast or brightfield training data, wrong organism (bacteria, Drosophila), 3D-only architecture, or wrong imaging modality (electron microscopy). The four remaining domain-compatible models were executed on all five test images via the model execution service and ranked by mAP.

#### Evaluation

Instance segmentation quality was assessed using mAP (mean average precision), computed as the mean F1 score over 18 IoU thresholds ranging from 0.1 to 0.95 in steps of 0.05^15^. At each threshold, predicted and ground-truth instances were matched greedily by descending IoU; matched pairs with IoU above the threshold were counted as true positives, remaining predictions as false positives, and unmatched ground-truth instances as false negatives. The mAP was computed as the mean F1 across all 18 thresholds, averaged over the five test images. Extending the threshold range to IoU = 0.1 captures partial overlaps that are invisible at the conventional IoU = 0.5 cutoff, providing a fuller picture of segmentation quality across all four models.

### COLLABORATIVE FINE-TUNING EXPERIMENT

#### Dataset

The PlantSeg lateral root dataset^5^ (SBIAD1392; 3D confocal microscopy of *Arabidopsis thaliana* lateral root, xylem pole pericycle nuclei) was used. Three volumetric movies (Movie 1–3) provided training data, each containing approximately 500 z-slices at full resolution. Per-movie p90 diameter normalisation factors (M1 × 0.3007, M2 × 0.3385, M3 × 0.4098) were computed to map nucleus diameters to the Cellpose-SAM training range (approximately 120 px). Images were pre-scaled accordingly and training was performed with `rescale=False`. In a real collaborative annotation session, annotators can measure representative cell diameters directly in the browser annotation tool, which is used to compute the rescaling factor automatically.

#### Test set

Three representative z-slices per movie were held out as the fixed evaluation set: Movie 1: z = 100, 250, 420; Movie 2: z = 80, 200, 360; Movie 3: z = 45, 110, 165 (9 slices total). Test images were rescaled using the same per-movie factors as the training data.

#### Annotation cycles

Four simulated collaborative annotation cycles were performed. Cycle 1 annotated 3 slices from Movie 1 (z = 20, 60, 150) for 100 epochs. Cycle 2 extended Movie 1 annotations to 7 slices (z = 20, 60, 150, 200, 310, 360, 460) for 500 epochs. Cycle 3 retained the same 7 Movie 1 slices and added 7 new slices from each of Movie 2 and Movie 3 (21 total) for 500 epochs. Cycle 4 added 7 further slices from each movie (42 total) for 500 epochs. New slices were chosen to cover complementary z-positions not already in the training set.

#### Model and training

Cellpose-SAM (Cellpose v4.0.7, `cpsam` checkpoint (ViT-L SAM encoder with convolutional readout)) was fine-tuned via the BioEngine Cellpose fine-tuning service (https://hypha.aicell.io/bioimage-io/services/cellpose-finetuning). Training hyperparameters were learning rate 1 × 10^− 6^ and weight decay 1 × 10^− 4^. Cycles 3 and 4 continued from the Cycle 2 checkpoint rather than restarting from the base model. Each cycle ran on a single NVIDIA A40 GPU, with 500 epochs requiring approximately 2.5 hours.

#### Evaluation

F1 score at IoU ≥ 0.5 was computed after each cycle using Hungarian matching. Per-movie averages across the three test slices are reported. Cumulative epoch counts were 0 (baseline), 100 (Cycle 1), 600 (Cycle 2), 1,100 (Cycle 3), and 1,600 (Cycle 4). The baseline (epoch 0) reflects Cellpose-SAM inference on rescaled test images with no fine-tuning. Mean F1 increased from 0.36 (baseline) to 0.71 (Cycle 4).

### EM MITOCHONDRIA ANALYSIS APPLICATION

The EM Mitochondria Analyzer (Fig. 2d) was built by an AI agent using the BioEngine SKILL.md to demonstrate application construction and deployment on BioEngine.

#### Dataset

The Lucchi++ FIB-SEM benchmark^11^ (mouse hippocampus CA1 region, 768 × 1024 px, 5 nm/px) was used. Ten evenly spaced test slices (z = 0, 18, 36, 54, 72, 90, 108, 126, 144, 162) with binary ground-truth mitochondria masks were evaluated.

#### Application architecture

The application (Hypha artifact ID at bioimage-io/fibsem-mito-analysis, browsable https://hypha.aicell.io/bioimage-io/artifacts/fibsem-mito-analysis) is a CPU-only Ray Serve deployment (2 CPUs, 0 GPUs, 4 GB RAM) that delegates all GPU inference to the model-runner service. The model-runner runs the `poisonous-spider` model (BioImage.IO Attention U-Net trained on the Lucchi++ training set^11^). No GPU is allocated to the application itself, illustrating how BioEngine applications can compose existing services without duplicating compute resources.

#### Inference pipeline

Input images are percentile-normalised (p1–p99). Each image (or, for large images, a series of overlapping 512 × 512 tiles) is passed to the model-runner service, which handles tiled inference and output stitching using the model’s specified block size. Overlapping tiles are blended using a Gaussian weight window before probability accumulation. The returned probability map is thresholded at 0.5, followed by morphological closing (disk radius 4) and removal of objects smaller than 300 px. A watershed transform seeded from distance-transform local maxima produces final instance masks.

#### Morphological profiling

Per-instance shape properties are computed from the labelled masks: area (µm^2^, converted using pixel size), aspect ratio (major axis divided by minor axis), eccentricity, and circularity. Instances with aspect ratio ≥ 2.0 are classified as elongated and those below as round.

#### Agent task

The AI agent received the BioEngine SKILL.md and a plain-language description of the desired analysis. Using the bundled Ray Serve deployment template and manifest example in the SKILL.md, the agent wrote the deployment script and manifest file, uploaded the package to the Hypha artifact store, and triggered deployment, producing a live endpoint at https://hypha.aicell.io/bioimage-io/services/fibsem-mito-analysis without manual steps.

