## Supplementary Figures for "BioEngine: scalable execution and adaptation of bioimage AI through agent-readable interfaces"

### Supplementary Information — BioEngine

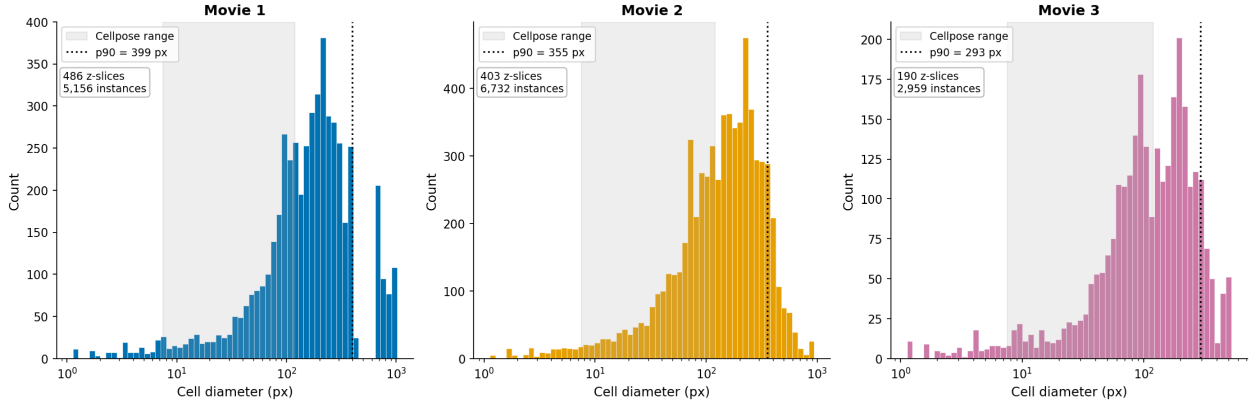


**Supplementary Fig. S1 | Nucleus size distributions in PlantSeg lateral root movies motivate per-movie diameter normalisation.** Histograms of equivalent nucleus diameter (log scale) for the three PlantSeg movies (Movie 1–3) used in the collaborative fine-tuning experiment. The grey region indicates the diameter range (7.5–120 px) on which Cellpose-SAM was trained. The dashed line marks the 90th-percentile (p90) diameter per movie (M1: 399 px, M2: 355 px, M3: 293 px). All three movies lie substantially outside the Cellpose-SAM training range, requiring per-movie rescaling to bring nuclei within the model's operating diameter window. Per-movie rescaling factors (M1 × 0.30, M2 × 0.34, M3 × 0.41) map p90 diameters to 120 px and were applied consistently to both training and test images.

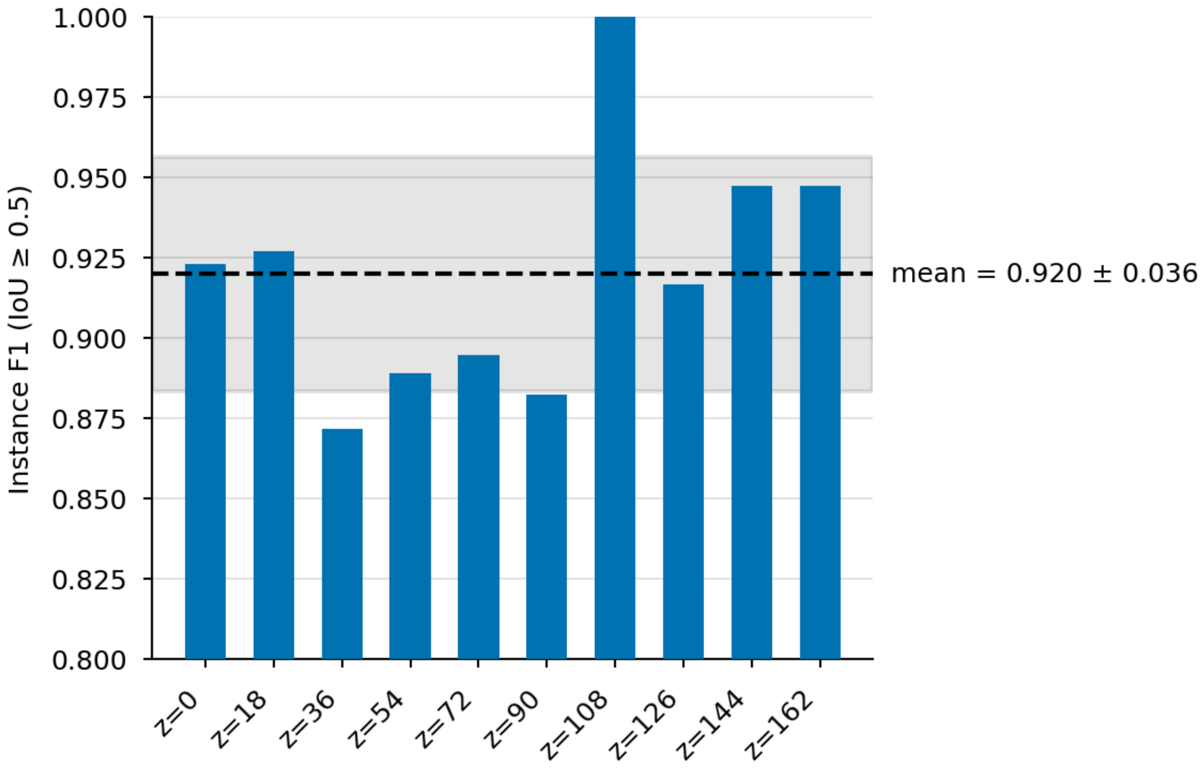


**Supplementary Fig. S2 | Quantitative validation of the EM Mitochondria Analyzer on the Lucchi++ benchmark.** Instance F1 at IoU ≥ 0.5 for the BioEngine-deployed poisonous-spider pipeline across 10 evenly spaced test slices of the Lucchi++ FIB-SEM dataset (neural tissue, 5 nm/px). The dashed line indicates the mean and the shaded band shows ± s.d. Mean F1 = 0.920 ± 0.037 (n = 10 slices).
